# Motion/direction-sensitive thalamic neurons project extensively to the middle layers of primary visual cortex

**DOI:** 10.1101/2021.07.07.451534

**Authors:** Jun Zhuang, Yun Wang, Naveen D. Ouellette, Emily Turschak, Rylan S. Larsen, Kevin T. Takasaki, Tanya L. Daigle, Bosiljka Tasic, Jack Waters, Hongkui Zeng, R. Clay Reid

**Affiliations:** Allen Institute for Brain Science, Seattle, WA, USA, 98109

## Abstract

The motion/direction-sensitive and location-sensitive neurons are two major functional types in mouse visual thalamus that project to the primary visual cortex (V1). It has been proposed that the motion/direction-sensitive neurons mainly target the superficial layers in V1, in contrast to the location-sensitive neurons which mainly target the middle layers. Here, by imaging calcium activities of motion/direction-sensitive and location-sensitive axons in V1, we find no evidence for these cell-type specific laminar biases at population level. Furthermore, using a novel approach to reconstruct single-axon structures with identified *in vivo* response types, we show that, at single-axon level, the motion/direction-sensitive axons have middle layer preferences and project more densely to the middle layers than the location-sensitive axons. Overall, our results demonstrate that Motion/direction-sensitive thalamic neurons project extensively to the middle layers of V1, challenging the current view of the thalamocortical organizations in the mouse visual system.

## Introduction

In mammalian visual systems, functionally specific thalamocortical projections from dorsal lateral geniculate nucleus (dLGN) to primary visual cortex (V1) serve as major feedforward inputs for cortical computation. When compared with higher mammals, the mouse dLGN shows more diverse response properties. Particularly, besides cells with classical spatial receptive fields that are sensitive to stimulus location (Grubb & Thompson, 2003; Denman et al., 2016), a significant portion of the cells in mouse dLGN are sensitive to motion direction (Marshel et al., 2012; Piscopo et al., 2013; Zhao et al., 2013; Durand et al., 2016; Suresh et al., 2016; Román Rosón et al., 2019). How these motion/direction-sensitive dLGN cells project to V1 is not fully understood. One current view is that the motion/direction-sensitive cells, resembling the W cells in cats, preferentially project to the superficial layers (layer 1), in contrast to the location-sensitive cells which have a middle layer bias (deep layer 3 and layer 4), resembling the X/Y cells in cats (Cruz-Martin et al., 2014, see review Seabrook et al., 2017). But the evidence for this laminar specificity is scarce and controversial: while one study found in support of this view that the middle layers in V1 receive slightly less direction-sensitive inputs from dLGN than the superficial layers (Kondo et al., 2016), another study found contradictory evidence that middle and superficial layers receive the same amount of direction-sensitive inputs from dLGN (Sun et al., 2016). This inconsistency is likely because both studies drew their conclusions from bulk labeled population distributions but neither investigated single axon morphology. In this study, we investigated the projection patterns of motion/direction-sensitive and location-sensitive dLGN axons in V1 at population level and, more importantly, at single-axon level. We found strong evidence that the motion/direction-sensitive dLGN axons project extensively to the middle layers in V1, providing an alternative model of functional specificity in mouse thalamocortical projects.

## Results

To measure the population distribution of different dLGN inputs to V1, we labeled dLGN cells and their axons with calcium indicator by injecting AAV viruses containing Cre-dependent GCaMP6s into the dLGN of Vipr2-IRES2-Cre-neo transgenic mice (Figure 1A), which have concentrated Cre expression in dLGN (Figure S1). We then measured the calcium activities of labeled dLGN axons in V1 in awake, head-fixed animals using two-photon imaging (Figure 1B, C). In total, 158 planes from 6 mice were imaged with imaging depths ranging from 50 to 400 μm below pia and calcium traces were extracted from 40008 regions of interest (ROIs) representing putative axonal boutons. We mapped the spatial receptive fields with locally sparse noise, and measured the tuning for orientation/direction, and spatial/temporal frequency (SF/TF) with full-field drifting gratings. A subset of experiments was carried out using adaptive optics in order to increase the fluorescence intensity and signal-to-noise ratio (Figure S2A, C). The response properties were statistically indistinguishable with and without adaptive optics (Figure S2B, D, E), so the data were pooled together.

**Figure 1.**
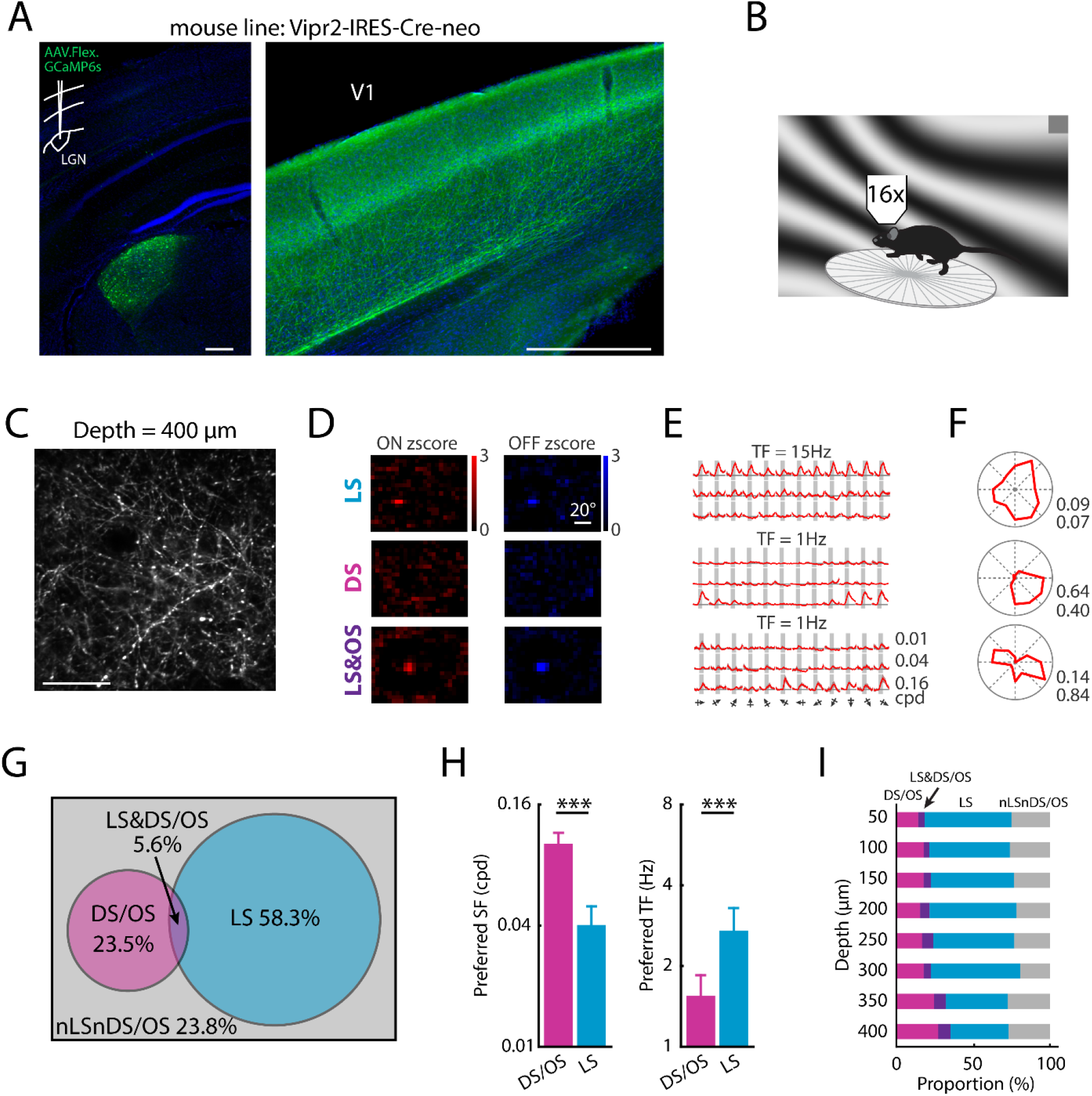
Labeling strategy, calcium imaging, classification, and depth distribution of dLGN boutons in V1. **(A)** GCaMP (green) expressions in a mouse prepared for calcium imaging experiments. Coronal sections. Inset: viral labeling. Blue: DAPI. Green: GCaMP6s. Scale bar: 500 μm **(B)** Sketch of *in vivo* imaging setup. **(C)** Mean projection of two-photon calcium images from an example imaging plane at 400 μm below pia. Scale bar: 50 μm. **(D-F)** The ON and OFF Z-score receptive fields, averaged calcium responses to gratings, and direction tuning curves, respectively, of three example putative boutons. Traces in **(E)**: mean df/f ± standard deviation across trials. Only responses to peak temporal frequency are shown. For each ROI, responses were normalized to peak responses. Grey boxes: time windows used to calculate responses. Columns: direction. Rows: spatial frequency. LS: location-sensitive. DS: direction-sensitive. OS: orientation-sensitive. Numbers in **(F)**, top: gDSI, bottom: gOSI. **(G)** Venn diagram describing the relations among the four different functional groups. **(H)** Comparisons of preferred spatial frequencies and temporal frequencies between DS/OS and LS boutons. The mean preferred SF and TF were calculated for each imaging plane and paired comparisons between DS/OS and LS boutons were made. ***: p < 0.001, Wilcoxon rank sum test. Bar graph: mean ± sem. **(I)** The ratio of each of the four groups in **(G)** across cortical depths.

Among all ROIs, 19967 were responsive to at least one type of displayed visual stimuli (Methods). Within these responsive boutons, many showed significant spatial receptive fields (RF strength ≥ 1.6, Method, Figure 1D) indicating sensitivity to stimulus locations (location-sensitive, LS), and many showed strong selectivity to one grating direction (gDSI > 0.5, direction-sensitive, DS, Figure 1E, F, middle) or two opposite grating directions (gOSI > 0.5, orientation-sensitive, OS, Figure 1E, F, bottom) indicating sensitivity to visual motion directions. The LS group (58.3%) and DS/OS group (23.5%) were largely separate with only 5.6% overlap (Figure 1G) which was significantly less than chance (0.583 × 0.235 = 13.7%, p=0.0000, Chi-square test). Consistent with previous electrophysiology studies (Piscopo et al., 2013), the DS/OS boutons preferred higher SF and lower TF than LS boutons (Figure 1H, DS/OS vs. LS, preferred SF: 0.10±0.02 vs. 0.04±0.01 cpd, p=1.6×10^−14^; preferred TF: 1.50±0.31 vs. 2.68± 0.61 Hz, p=3.3×10^−14^, Wilcoxon rank test). A small number of boutons showed distinct types of response properties such as suppressed-by-contrast (data not shown) as reported previously (Piscopo et al, 2013, Durand et al., 2016, Suresh et al., 2016). But since they were relatively rare and not the focus of this study, they were grouped into the non-LS and non-DS/OS (nLSnDS/OS) group. Both LS and DS/OS boutons distributed evenly across all measured cortical depths (Figure 1I), consistent with one previous report (Sun et al., 2016). In addition, all measured response metrics showed no depth bias except preferred SF, which was higher at greater depths (Figure S3). Thus, LS and DS/OS boutons appear to form two distinct feedforward input types in V1 that are present in both superficial (layer 1 and superficial layer 2/3) and middle layers (deep layer 2/3 and layer 4).

In analyzing a population, it is important to link the boutons that arise from the same axon, which we achieved by using a correlation-based hierarchical clustering procedure (Methods, Liang et al., 2018). With this procedure, ROIs with highly correlated activities were grouped into separate clusters (Figure 2A, B). and the correlation coefficients showed non-overlapping distribution between within-cluster and between-cluster bouton pairs (Figure 2F). The boutons from single clusters showed highly correlated calcium activities (Figure 2C) and near-identical response properties (Figure 2D, E), further confirming their high likelihood of being from same axons. For each cluster, we used summed calcium traces weighted by ROI intensities of its boutons to calculate its response properties and used same criteria to group them into DS/OS clusters and LS clusters.

**Figure 2.**
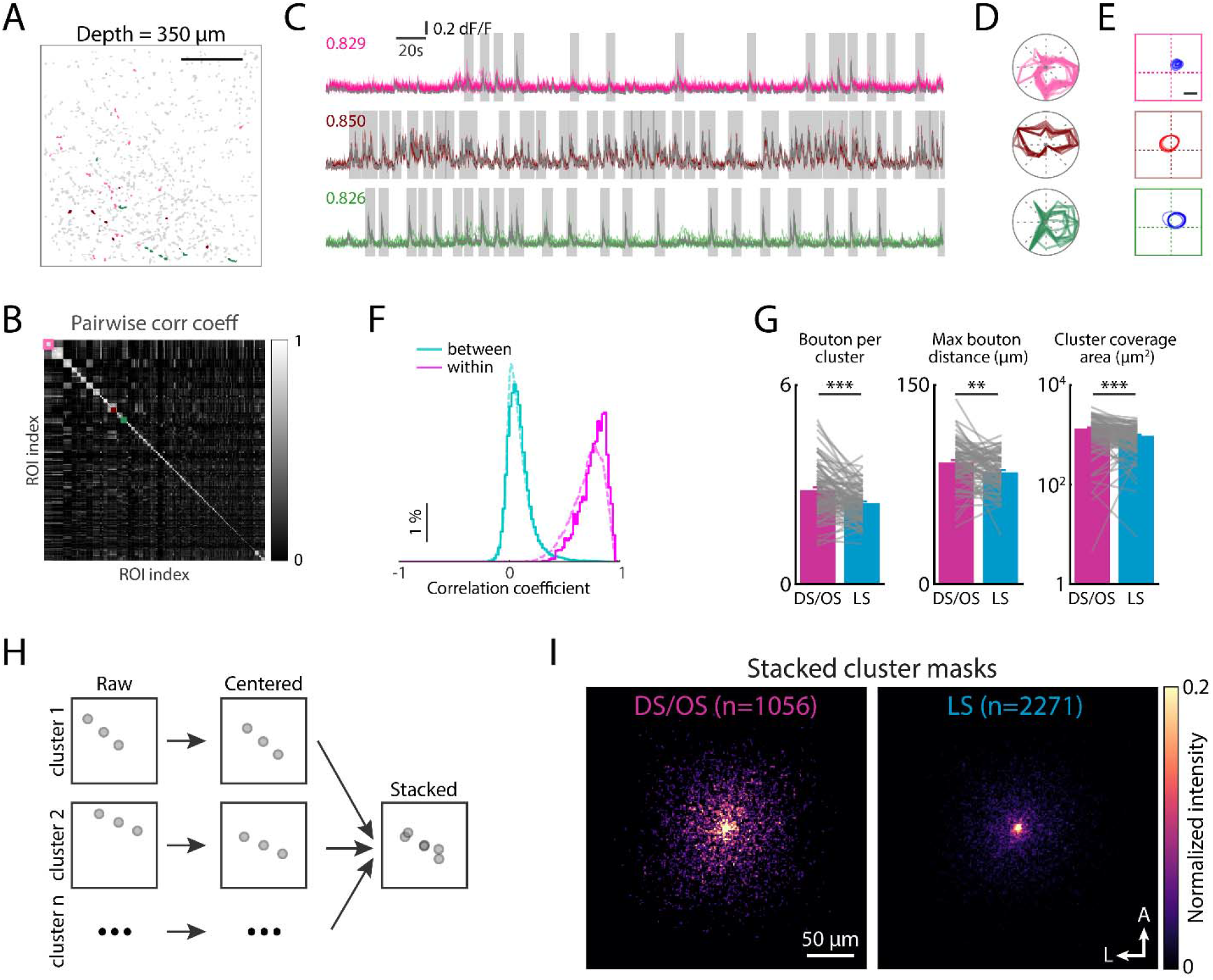
DS/OS bouton clusters have larger bouton spread than LS bouton clusters **(A)** An example imaging plane showing all active boutons (grey) and three example bouton clusters (colored). Scale bar: 50 μm **(B)** Clustered correlation coefficient matrix for the imaging plane in **(A)**. Example clusters in **(A)** are indicated by colored boxes. **(C)** Calcium traces from the example clusters in **(A)**. Each row represents a cluster with matching color. Colored traces: traces from each bouton in the cluster. Grey traces: weighted average traces for each cluster. Numbers: mean pair-wise correlation coefficient. Grey boxes: time windows of detected calcium events used for performing correlation (Methods). **(D, E)** Superimposed direction tuning curves and receptive field contours, respectively, of individual boutons from each cluster. **(F)** Normalized distribution of correlation coefficients between within-cluster boutons and cross-cluster boutons. Solid line: data from the example plane in **(A)**. Dashed line: data from all imaging planes. **(G)** Comparisons of bouton per cluster, maximum bouton distance, and cluster coverage area between DS/OS and LS clusters. Means of each matric were calculated for each plane and paired comparisons were made between DS/OS and LS clusters. Each grey line represents one imaging plane. **: p< 0.01. ***: p<0.001. Wilcoxon rand sum test. **(H)** Stacked masks are generated by superimposing centered individual cluster masks. **(I)** Stacked masks showing DS/OS clusters having larger bouton spread than LS clusters. A: anterior; L: lateral.

We then compared the structural properties between the DS/OS and LS clusters. For each cluster, we measured the bouton number, maximum bouton distance, and cortical coverage area. Interestingly, the values for DS/OS clusters were greater than for LS clusters for all three metrics (Figure 2G, DS/OS vs LS, bouton number: 2.84±0.08 vs. 2.44±0.05, p=7.2×10^−6^; max bouton distance: 91.5±2.0 vs. 84.3±1.6 μm, p=0.002; axon coverage: 1324±81 vs. 957±51 μm^2^, p=3.8×10^−6^, Wilcoxon rank test). To directly visualize the bouton spread of DS/OS and LS clusters, we generated stacked population cluster masks for each group (Figure 2H). The results showed that the DS/OS clusters had substantially greater bouton spread than the LS clusters consistent with above measurements (Figure 2I, distances from every bouton to its cluster center, DS/OS vs. LS, 42.2±27.3 μm vs. 39.7 ± 27.9 μm, p=1.22×10^−8^, Mann-Whitney test). The greater bouton count and larger coverage of DS/OS cluster suggests that the DS/OS axons had denser axon arbors than LS axons.

Using *in vivo* two-photon images to estimate axon structures, however, had substantial incomplete sampling: a two-photon imaging plane (~180 × 180 × 6 μm) only sampled less than 1% of an axon arbor which can extend to a volume of ~500 × 500 × 500 μm (Antonini et al., 1999, Figure 4). To overcome this limitation, we developed a novel approach that allowed us to investigate the structures of complete axon arbors with identified *in vivo* response properties (Figure 3A, Methods). In this approach, dLGN axons were sparsely labeled with GCaMP6s and blood vessels were labeled with a red fluorescent dye (Dextran Texas Red). The same visual stimuli and imaging protocols were used to identify response types. The brain tissue was then fixed and sectioned tangentially, followed by antibody staining to enhance GCaMP signal and a counterstain to label blood vessels. Finally, the tissue was cleared using the CUBIC clearing method (Susaki et al., 2015). The labeled and cleared tissue volumes were then imaged by a confocal microscope. Using the labeled blood vessels, the *in vivo* two-photon images were spatially aligned to the confocal image stacks (Figure S4), allowing precise coregistration between the functional recordings of the boutons and their anatomical locations allowing tracing and 3D reconstruction. Due to the high sparsity of the labelling, different boutons with very similar response properties (even across different imaging sessions) always appeared to belong to same reconstructed axons (Figure 3B-D), confirming the reliability of our coregistration and the completeness of our reconstructions.

**Figure 3.**
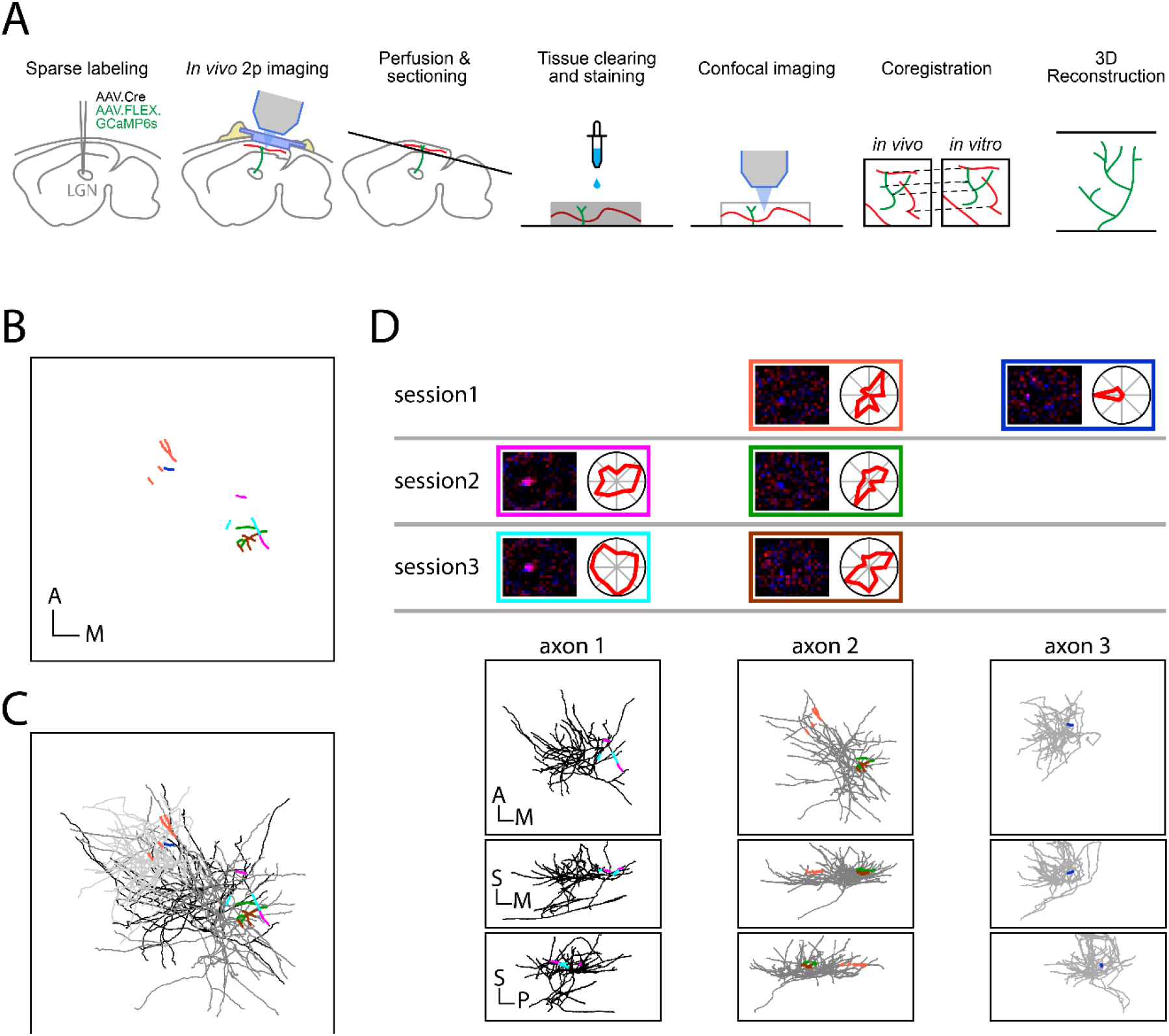
Reconstructing 3D structures of axons with identified response properties. **(A)** sketches showing the workflow to reconstruct 3D structures of axons with identified *in vivo* response properties. Mouse brain was shown as sagittal sections. dLGN axons was sparsely labeled by injecting mixture of AAV9-hSyn-Cre and AAV1-Syn-FLEX-GCaMP6s (1:40000) into the dLGN of wildtype mice. Blood vessels were labeled by fluorescent dyes in both *in vivo* and *in vitro* imaging experiments and used as fiducials for coregistration across imaging modalities. Green lines: labeled axons. Red lines: labeled blood vessels. **(B)** All axon segments from a single mouse with identified response properties (pooled from three imaging sessions). Each color represents a bouton cluster identified using calcium activity correlation-based clustering method (Figure 2). In total, 6 axon segments were identified. **(C)** Three reconstructed axon arbors (marked as different grey levels) contained the 6 segments marked in **(B)**. **(D)** Top, the receptive fields and direction tuning curves from the 6 segments marked in **(B)** with color matched boxes. Note the segments imaged in different sessions can show similar (within column) or different (cross column) response properties. Bottom, three views of the three reconstructed axon arbors showing that the segments having similar response properties across different imaging sessions belong to same arbors and segments having different response properties belong to different arbors.

**Figure 4.**
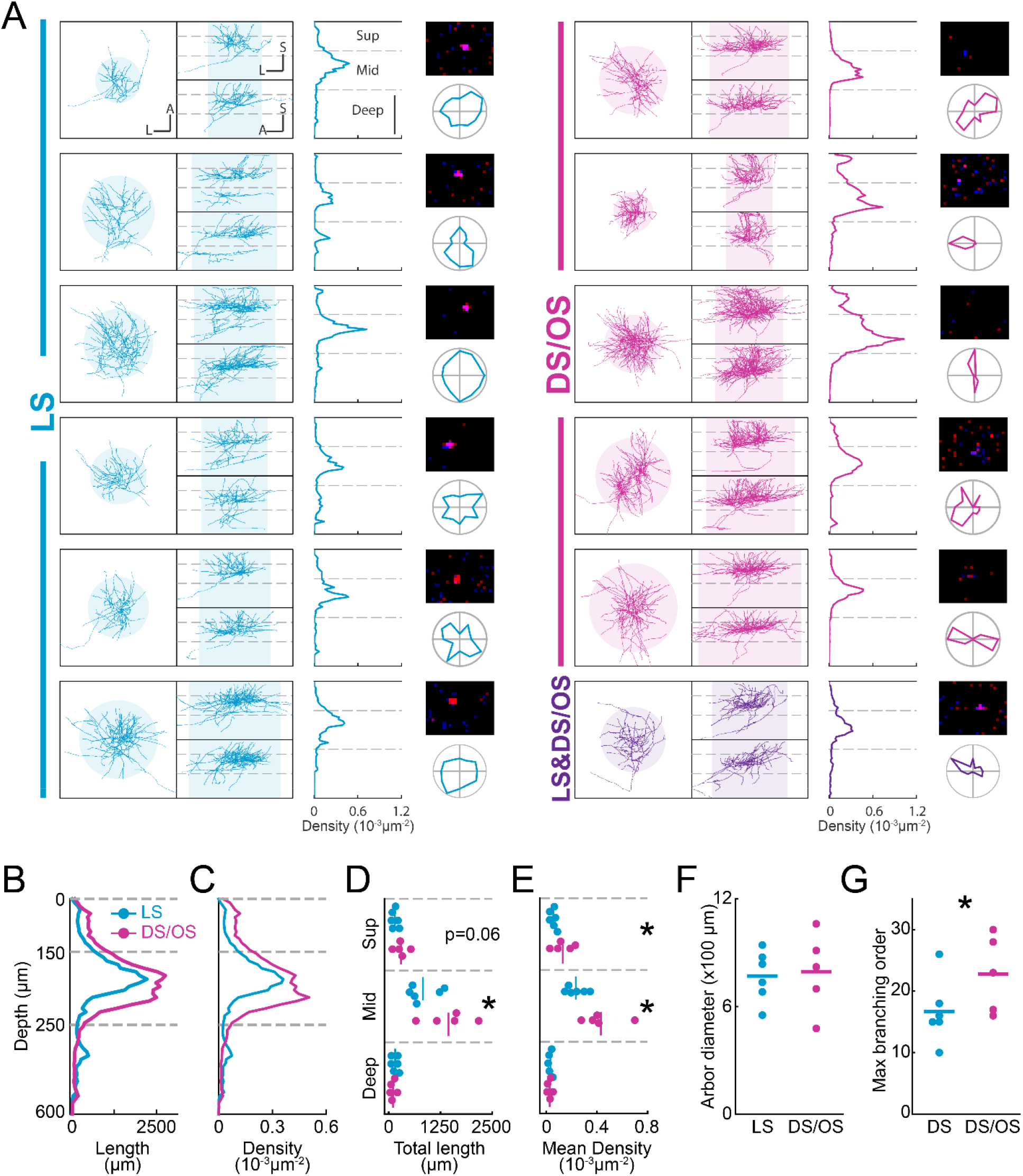
DS/OS axons have denser projection in superficial and middle layers than LS axons. **(A)** 3D structures and response properties of all reconstructed axons. For each axon, the three views (left), density across depth (middle), receptive field (top right), and direction tuning curve (bottom right) are shown. Scale bar: 200 μm. A: anterior, L: lateral; S: superficial. Horizontal dashed lines: borders to define superficial (0 – 150 μm), middle (150 – 250 μm), and deep (250 – 600 μm). Shaded circles: horizontal arbor extent defined as a circle extending from center of mass and encompassing 95% of all segments. Shaded boxes: volumes by extruding the arbor extents along cortical depth for calculating arbor density. Numbers, indices for each axon. **(B, C)** Comparisons of mean total length and mean density, respectively, across cortical depth between DS/OS and LS axons. Horizontal dashed lines match those in **(A)**. **(D, E)** Comparisons of total length and density, respectively, between DS/OS and LS axons in superficial, middle, and deep layers (separated by horizontal dashed lines). Colored bars: measurements from single axons ordered by the indices in **(A)**. Open black circle: mean for each group at each depth. *: p < 0.05. Mann-Whitney U test. **(F, G)** Comparisons of arbor diameter and maximum branching order between DS/OS and LS axons. Colored bars: measurements from single axons ordered by the indices in **(A)**. Open black circle: mean. *: p<0.05. Mann-Whitney U test.

In total, six LS, five DS/OS, and one LS&DS/OS axons were reconstructed (Figure 4A). All axons projected extensively to the middle layers (150 – 250 μm below pia) and extended to the superficial layers (0 – 150 μm below pia). Three LS axons (axon 2, 4, and 5 in Figure 4A) but no DS/OS axons showed secondary clusters in the deep layers (250 – 600 μm below pia). Although no significant difference was found in the horizontal coverages (shaded circles in Figure 4A, Methods) between LS and DS/OS axons (Figure 4F, coverage diameter, DS/OS vs. LS, 794 ± 221 vs. 770 ± 142 μm, p=0.46, Wilcoxon rank test), there were prominent differences in segment length and arbor density (defined as segment length divided by a cylindrical volume encompassing the axon, Method) between superficial and deep layers and between DS/OS and LS axons. First, both DS/OS and LS axons showed greater segment length and arbor density in the middle layers than in the superficial layers (Figure 3B-E, middle vs superficial, DS/OS length: 28.9 ± 11.3 vs. 5.9 ± 3.2 mm, p=0.03; LS length: 16.5 ± 7.3 vs. 3.0 ± 1.6 mm, p=0.02; DS/OS density: 43.1 ± 15.9 vs. 13.0 ± 8.0 × 10^−5^ μm^−2^, p=0.03; LS density: 23.4 ± 7.7 vs. 5.7 ± 2.5 × 10^−5^ μm^−2^, p=0.02, Wilcoxon test). In addition, when compared with the LS axons, the DS/OS axons had greater segment length and arbor density in the middle layers (Figure 3B-E, length: p=0.04; density: p=0.01, Mann-Whitney test). This difference was also apparent in the superficial layers but only the difference in arbor density reached statistical significance (length: p=0.06; density: 0.03, Mann-Whitney test). No significant difference was found in the deep layers (length: p=0.31; density: p=0.51, Mann-Whitney test). Finally, the DS/OS arbors showed higher maximum branching order than the LS axons consistent with their higher arbor density (Figure 3G, DS/OS vs LS, 22.8 ± 6.3 vs. 16.7 ± 5.3, p=0.049, Mann-Whitney test). These results indicated that DS/OS axons, like the LS axons, preferentially target the middle layers, but with denser axon arbors.

## Discussion

In this study, we show strong evidence, at both population and single-cell level, that the motion/direction-sensitive dLGN neurons project extensively to the middle layers in V1, arguing against the hypothesis predicting their superficial projection bias (Seabrook et al., 2017), but consistent with the findings from other species (Hei et al., 2015; Bereshpolova et al., 2019). These results suggest that the motion sensitivity in the V1 middle-layer cells can be partially inherited from the dLGN motion/direction-sensitive inputs, although it can also be constructed from non-motion-sensitive inputs (Lien & Scanziani, 2018). Our results also argue against the proposed homology between the DS/OS pathway in mice and the W-pathway in cats (Seabrook et al., 2017). The DS/OS cells in this study project heavily to middle layers (Figure 4) and preferred low TF and high SF than LS cells (Figure 1H, Piscopo et al., 2013), whereas the W cells in cat dLGN predominantly project to the superficial layers (Anderson et al., 2009) and prefer lower SF than X- and Y-cells (Sur & Sherman, 1982).

Compared with the rapid progress in mapping complete local functional connectomes in mouse V1 (microns-explorer.org), our knowledge about how V1 receives its major inputs from dLGN is surprisingly incomplete. A recent large-scale survey has provided morphological classifications of thalamocortical projections (Peng et al., 2020), but, without *in vivo* physiology, the insights it can provide for cortical computation are indirect. Our study, with its unique strength in linking *in vivo* functions and single-axon morphologies, will complement these large-scale studies by providing function-structure correspondence of the major feedforward inputs to V1.

## Methods

### Surgery and animal preparation

For the dense labeling experiments, a 1:3 mixture of AAV1-CAG-mRuby3 (custom made from plasmid Addgene 107744, titer: 1.6×10^12^ vg/ml) and AAV1-Syn (or CAG)-FLEX-GCaMP6s (Addgene: 100845-AAV1, titer 4.3×10^13^ vg/ml or 100842-AAV1, titer 1.8×10^13^ vg/ml, respectively) was injected into the dLGN of six Vipr2-IRES2-Cre-neo mice (3 male, 3 female, 200nL each). For the sparse labeling experiments, a 1:1 mixture of AAV9-hSyn-Cre (1:40000 dilution, Addgene: 105553-AAV9, titer: 3.3×10^13^vg/ml) and AAV1-Syn-FLEX-GCaMP6s (Addgene: 100845-AAV1, title: 2.5×10^13^vg/ml) was delivered into the dLGN of 6 wild type C57BL/6J mice (3 male, 3 female, 100nL each, Economo et al., 2016). Briefly, injection pipette was slowly lowered into left dLGN (2.3 mm posterior 2.3 mm lateral from bregma, 2.6 mm below pia) through a burr hole on the skull. 5 minutes after reaching the targeted location, the virus mixture was injected into the brain over 10 minutes by a hydraulic nanoliter injection system (Nanoject III, Drummond). The pipette then stayed for an additional 10 minutes before it was slowly retracted out of the brain. Immediately after injection, a titanium head-plate and a 5 mm glass cranial window were implanted over left V1 (Goldey et al., 2014) allowing *in vivo* two-photon imaging during head fixation.

After surgery, the animals were allowed to recover for at least 5 days before retinotopic mapping with intrinsic signal during anesthesia (Juavinett et al., 2017). After retinotopic mapping, animals were handled and habituated to the imaging rig for two additional weeks (de Vries et al., 2020) before *in vivo* two-photon imaging.

All experiments and procedures were approved by the Allen Institute Animal Care and Use Committee.

### *In vivo* two-photon imaging

In awake animals, the calcium activities were recorded with a conventional two-photon microscope or with a multi-plane two-photon microscope (described below). In both microscopes, a 16x/0.8 NA water immersion objective (Nikon 16XLWD-PF) was rotated to 24 degrees from horizontal to image visual cortex using a commercial rotating head (Sutter MOM). Emitted light was first split by a 735 nm dichroic mirror (FF735-DiO1, Semrock). The short-wavelength light was filtered by a 750 nm short-pass filter (FESH0750, Thorlabs) and a 470-588 nm bandpass emission filter (FF01-514/44-25, Semrock) before collected as GCaMP signal, while the long-wavelength light was filtered by a 538-722 nm band-pass emission filter (FF01-630/92-30, Semrock) before collected (mRuby for dense labeling experiments and Dextran-Texas Red for sparse labeling experiments). Image acquisition was controlled using Vidrio ScanImage software for both scopes (Pologruto et al., 2003, Vidrio LLC). To maintain constant immersion of the objective, we used gel immersion (Genteal Gel, Alcon).

For experiment with dense labeling (6 mice), thalamocortical axons in V1 were imaged at 8 cortical depths (50, 100, 150, 200, 250, 300, 350, and 400 μm below pia). Imaging was done in a columnar fashion: at each cortical location calcium activities at 3-8 depths were imaged plane-by-plane over multiple sessions.

For experiments with sparse labeling/axon reconstruction (6 mice), additional 30-60 uL of Dextran Texas Red (Thermo Fisher, D3328, 25 mg/mL solution with saline) was injected subcutaneously ~20 minutes before single-plane two-photon imaging sessions to label blood vessels. For each imaging session, a local z-stack (field of view 358 × 358 μm from pia to depth of 500 μm with 4 μm step) was recorded to aid coregistration.

### Single-plane two-photon imaging

Two-photon excitation was generated by laser illumination from a Ti:sapphire laser (Coherent Chameleon Ultra II) tuned to 920 nm. A single z-plane (179.2 × 179.2 μm) was imaged for each session at a frame rate of about 30 Hz with an 8 KHz resonate scanner (Cambridge Technology, CRS 8K).To maintain constant imaging depth automatic z-drift correction functions were implemented for experiments using the MOM motors. Briefly, a correction z-stack (± 50 μm from targeted depth, 2 μm step depth) was recorded before each imaging session and, during the session, the current imaging plane was continuously compared to each plane in the correction z-stack. If a drift in depth was detected, the stage was automatically adjusted to compensate for the drift, thus maintaining constant imaging depth. We found this procedure crucial to our experiments since the boutons are small objects and a few-micron-drift in depth would result in imaging a different set of boutons.

### Multi-plane two-photon imaging

In this custom-built, multi-plane two-photon microscope (DeepScope, Liu et al., 2018), a liquid crystal spatial light modulator (SLM; HSP-512, Meadowlark Optics) shapes the pupil wavefront to implement fast-focusing and adaptive optics. The objective pupil was slightly underfilled (~0.65 effective NA) and correction of systemic aberrations was performed with fluorescent beads to maintain near diffraction-limited focusing over a 200 um range. Two-photon excitation was produced by laser light from a commercial solid-state laser (Spectra-Physic Insight X3 laser) tuned to 940 nm. With this microscope, we simultaneously recorded calcium activity from planes at 5 different depths (50, 100, 150, 200, 250 μm) in single imaging sessions. Individual frames (125 × 125 μm with 512 × 512 pixels resolution) were acquired at an overall framerate of ~37 Hz with a volume rate of 7.4 Hz. The DeepScope showed nearly zero z-drift for a prolonged duration (< 2 μm over 24 hours), so we did not implement z-correction in sessions using DeepScope.

In another set of experiments, we used DeepScope to assess the effect of adaptive optics adjusted on individual animals. The correction procedure was similar to the method described in Sun et al., 2016. For two mice, 1 μm beads (Thermo Fisher, F8821) were deposited on top of the brain surface under the coverglass during the initial surgery. Prior to the imaging session, modal optimization over 12 Zernike modes (up to j = 15 Noll ordering, excluding piston and tilt) was run to identify the SLM pattern that maximized the beads’ fluorescent signal. Then, during the imaging session, this SLM pattern was put on and off alternatively for consecutive two-photon imaging frames (one frame on, one frame off, repeated) and drifting gratings were displayed. After imaging, the interleaving movie was separated into two movies: one with adaptive optics and the other without. Bouton’s tuning properties were then extracted from each movie and compared against each other.

All imaging sessions were performed during head fixation with the standard Allen Institue Brain Observatory *in vivo* imaging stage (de Vries et al., 2020).

### Visual stimulation

All visual stimuli were generated and displayed by Retinotopic_Mapping python package (https://github.com/zhuangjun1981/retinotopic_mapping, Zhuang et al., 2017) over PsychoPy software (https://www.psychopy.org, Peirce et al., 2019) on a 24-inch LCD monitor (ASUS PA248Q, frame rate 60 Hz, 1920 × 1200 pixels, mean luminance 45.3 cd/m^2^) placed 15 cm from the mouse’s right eye (covering 120° × 95° of monocular visual space). We displayed locally sparse noise and full-field drifting grating in each imaging session to measure receptive fields and orientation/direction/spatial and temporal frequency tuning properties, respectively. In most sessions, we also displayed a five-minute full-field mid-luminance gray to measure spontaneous activity. For locally sparse noise, bright and dark squares (5° × 5°) were displayed in a random sequence on a grid tiling the entire monitor. At any given time, multiple squares could be displayed but the minimum distance between those squares should be no less than 50°. Each square lasted 100 ms and in total was displayed 50 times. For drifting gratings, the combinations of 12 directions (every 30°), 3 spatial frequencies (0.01, 0.04 and 0.16 cpd) and 3 temporal frequencies (1, 4, 15 Hz) were displayed. Each display lasted 1 second and was spaced by 1-second mean luminance grey period. In total, 3 × 3 × 12 + 1 (blank) = 109 conditions were randomly displayed in each iteration and the whole sequence contained 13 iterations. Although the SFs and TFs presented in our stimulus set were not enough to map comprehensive tuning curves (limited by total imaging time of each session), they were enough to be used for comparisons between different functional groups. All stimuli were spherically corrected so that they were presented with accurate visual angles on the flat screen (Zhuang et al., 2017).

### two-photon Image preprocessing

The recorded two-photon movies for each imaging plane were first temporally averaged across every 5 frames (Sutter scope) or 2 frames (DeepScope) retaining an effective temporal frequency of ~6 Hz. Motion-corrected was then performed on the red channel (mRuby in densely labeled samples and Texas Red in sparsely labeled samples) using rigid body transform based on phase correlation by a custom-written python package (https://github.com/zhuangjun1981/stia, Zhuang et al., 2017). The resulting correction offsets were then applied to the green channel (GCaMP). To generate regions of interest (ROIs), the motion-corrected movies were further temporally downsampled by a factor of 3 and then processed with constrained non-negative matrix factorization (CNMF, Pnevmatikakis et al., 2016) implemented in the CaImAn python library (https://github.com/flatironinstitute/CaImAn, Giovannucci et al., 2018). The resulting ROIs were filtered by their size ([1.225, 12.25] μm^2^), position (ROIs within the motion artifacts were excluded) and overlap (for ROIs with more than 20% overlap, the smaller ones were excluded). For each retained ROI, a neuropil ROI was created as the region between two contours by dilating the ROI’s outer border by 1 and 8 pixels excluding the pixels within the union of all ROIs. Same procedures of neuropil subtraction in our previous studies (Zhuang et al., 2017; de Vries et al., 2020) was then applied. As reported previously, a high skewness is an indication of active calcium activity (Mukamel et al., 2009, Dipoppa et al, 2018). Only the ROIs with skewness greater than 0.6 were defined as “active” boutons and were included in this study. For each imaging session, a comprehensive file in Neurodata Without Borders (nwb) 1.0 format was generated to store and share metadata, visual stimuli, all preprocessing results, and final calcium traces using the “ainwb” package (https://github.com/AllenInstitute/nwb-api).

### Bouton clustering

The correlation-based bouton clustering procedure was based on previously reported algorithms (Petreanu et al., 2012; Liang et al., 2018) and the same procedure was performed on each imaging plane. First, for each ROI, calcium events were detected as up-crosses over a threshold of 3 standard deviations above the mean in its calcium trace (gaussian filtered with a sigma of 0.1 sec). A period of 3 seconds before and after each onset was defined as an event window and the union of all event windows for a particular ROI was saved. Second, for any given pair of boutons, the union of event windows from both boutons was generated and the calcium trace within the union window was extracted and concatenated for each bouton. Then a Pearson correlation coefficient of the two concatenated traces was calculated for this pair. By performing this procedure on all pairs of active ROIs, we generated a correlation coefficient matrix for each imaging plane. We found the event detection important because it confined the correlation to the period in which at least one bouton in the pair was active thus avoided correlating the noise during the inactive period. Third, the correlation coefficient matrices were further thresholded to reduce noise: the correlation coefficients for a given bouton were maintained if the coefficients were larger than 0.5 or if they exceeded 3 standard deviations above the mean value of all the coefficients between this bouton and all others. Otherwise, they will be set to 0. Third, a hierarchy clustering was performed to a given imaging plane using “1 – thresholded correlation coefficient matrix” as the distance matrix using Scipy.cluster.hierarchy library with a “weighted” method (https://docs.scipy.org/doc/scipy/reference/generated/scipy.cluster.hierarchy.linkage.html). Fourth, we use a threshold of 1.3 to separate clusters since ~ 1.5 shows up as a relatively natural cut-off in the dendrograms. This threshold appeared somewhat conservative on the clustered correlation coefficient matrix. Fifth, calcium trace of each bouton cluster was calculated as mean calcium trace of all boutons belonging to this cluster, weighted by the sum of their ROI masks.

For each bouton cluster, three values were extracted to estimate the morphology: (1) bouton number of this cluster; (2) maximum distance among all bouton pairs belonging to this cluster; (3) area of the convex polygon encapsulated by all the boutons belonging to this cluster. For axons with only one bouton, the metrics 2 and 3 were set to be “nan” and were excluded from statistical analysis.

To generate stacked cluster mask, we first selected bouton clusters with at least two boutons for each group. Then the binary mask of each individual cluster was centered to its own center of mass. All centered masks were summed together to generate a summed population mask for each group, normalized by its peak. LS&DS/OS bouton clusters were excluded from this analysis to avoid double counting.

### Spatial receptive field analysis

We calculated the units’ spatial receptive fields from their responses to the locally sparse noise stimulus using reverse correlation analysis (Zhuang et al., 2013, 2017). For each stimulus location, the df/f value was calculated as (response – baseline) / baseline (with mean calcium trace [0, 0.5] second after stimulus onset as response and [−0.5, 0] second before onset as baseline). From this df/f amplitude map, a z-score map was calculated by subtracting the mean and dividing the standard deviation of the entire map. The z-score map was then smoothed (Gaussion filter, sigma = 1 pixel) and up-sampled by a ratio of 10 with cubical interpolation. RF strength was defined as the peak value of this z-score map. ROIs with a RF strength no less than 1.6 were defined as having a significant spatial RF. For each significant spatial RF, an RF mask was generated by thresholding either with a value of 1.6 (for maps with a peak less than 4) or with a value of 40% of its peak (for maps with a peak greater than 4).

### Grating response analysis

The units’ responses to drifting gratings were analyzed by the event-triggered average procedure similar to the RF response analysis (0.5 second before onset as baseline and 1 second after as response). We used same inclusion criteria used in our previous large-scale study (de Vries et al., 2020). The direction tuning curve was extracted at peak TF/SF conditions and, if the minimum response from this curve was below zero, an offset was added to the whole curve so that the minimum was zero. The SF and TF tuning curves were extracted by a similar procedure.

We calculated global direction selectivity index (gDSI) as

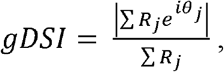

global orientation selectivity index (gOSI) as

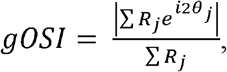

and the preferred direction as the angle of Σ *R*_*j*_*e*^*i*θ_*j*_^,. Where j represents different direction conditions, R represents df/f response in each direction and θ represents the direction. We defined a unit to be orientation-selective if either its gOSI was greater than 0.5 and a unit to be direction-selective if its gDSI was greater than 0.5.

From the SF tuning curve, we calculated the preferred SF as

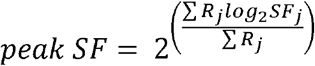

Where j represents different SF conditions. From TF tuning curve, we calculated the preferred TF as

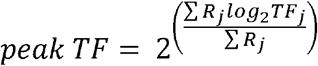

Where j represents different TF conditions.

### Perfusion

To histology experiment, the brain tissues were collected by transcardial perfusion. Briefly, mice were anesthetized with 5% isoflurane and 10 ml of saline (0.9% NaCl) followed by 50 ml of freshly prepared 4% paraformaldehyde (PFA) was pumped intracardially at a flow rate of 9 ml/min. Brains were immediately dissected and post-fixed in 4% PFA at room temperature for 3-6 hours and then overnight at 4 °C. After fixation, brains were incubated in PBS with 10% sucrose and then stored in PBS with 30% sucrose until sectioning.

### Histology for expression characterization

To characterize the Cre expression pattern, Vipr2-IRES2-Cre-neo mice were crossed with the Ai14 reporter line (Madisen et al., 2010) to generate Vipr2-IRES2-Cre-neo/wt; Ai14/wt animals. The brains of these animals were cut into 50 μm sections by a freezing-sliding microtome (Leica SM 2101R). Sections with dLGN and V1 were then mounted on gelatin-coated slides and cover-slipped with mounting media (Prolong Diamond Antifade Mounting Media, P36965, ThermoFisher).

To verify the injection location and GCaMP expression, the brain tissues from mice in dense labeling experiments were collected and sectioned with a similar procedure after all imaging sessions with additional steps of antibody staining to enhance the GCaMP signal before mounting and coverslipping. During antibody staining, sections containing dLGN and V1 were blocked with 5% normal donkey serum and 0.2% Triton X-100 in PBS for one hour, incubated in an anti-GFP primary antibody (1:5000 diluted in the blocking solution, Abcam, Ab13970) for 48-72 hours at 4 °C, washed the following day in 0.2% Triton X-100 in PBS and incubated in a Alexa-488 conjugated secondary antibody (1:500, 703-545-155, Jackson ImmunoResearch) and DAPI.

The sections were then imaged with Zeiss AxioImager M2 widefield microscope with a 10x/0.3 NA objective. Fluorescence from antibody enhanced GCaMP and mRuby3 were extracted from filter sets Semrock GFP-1828A (excitation 482/18 nm, emission 520/28 nm, dichroic cutoff 495 nm) and Zeiss # 20 (excitation 546/12 nm, emission 608/32 nm, dichroic cutoff 560 nm), respectively.

### Histology for axon reconstruction

To reconstruct the sparsely labeled dLGN axons, brain sections from mice in the sparse labeling experiments were cut tangentially to the imaging window surface with thickness of 350 or 400 μm on a vibrotome (Leica VT1000). The sections were then (1) incubated in PBS with 8% SDS for 48 hours at 37°C for initial clearing; (2) blocked by blocking solution NDSTU (5% Normal Donkey Serum, 4M Urea, 0.2% Triton X-100) at room temperature for 1 hour; (3) incubated in primary antibody for GFP (1:5000 in NDSTU, Abcam, Ab13970) at room temperature for 48 hours; (4) incubated in secondary antibody (1:500 703-545-155, Jackson ImmunoResearch) and lectin (for labeling blood vessels, 2 μg/mL, Vector Laboratories, DL1178) at room temperature for 48 hours; and (5) mounted with CUBIC tissue clearing solution (urea 25 wt%, Quadrol 25 wt%, Triton X-100 15 wt% in dH_2_O, Susaki et al., 2015) with spacers of appropriate depth (SunJin Lab, iSpacer).

The relevant regions (field of view 1.7 × 1.7 mm or 1.7 × 1.2 mm) were then imaged by a confocal microscope (Olympus FV3000) as tiled z-stacks (30x oil immersion objective with resolution 0.414 (x) × 0.414 (y) × 0.5 (z) μm, excitation 488 nm / emission [500, 540] nm for axons and excitation 640 nm / emission [650, 750] nm for blood vessels).

### Coregistration, reconstruction, and morphology analysis

Tiled confocal stacks were first stitched by TeraStitcher (https://abria.github.io/TeraStitcher/, Bria et al., 2012). The stitched volumes were rotated to match the standard orientation (up: anterior, left: lateral). Using surface vasculature, the field of view of each 2p session from the same mouse was located in the confocal volume. By carefully following the descending blood vessels in both 2p and confocal volumes in the red channel, the imaged depths were reached, and the imaged axon segments were identified in the green channel. From those identified axon segments, the complete axon arbor was manually traced using TeraFly/Vaa3d software (https://alleninstitute.org/what-we-do/brain-science/research/products-tools/vaa3d/).

To quantify the morphological features, we calculated the total length, maximum branching number, 2D diameter, and density for each reconstrued axon arbor. To calculate 2D diameter, we first collapse all segments into a 2D xy plane, then the dimeter of a circle centered at the center of mass and encompasses 95% of all segments was defined as the axon’s 2D diameter. The density was defined as the total length divided by the volume of the cylinder constructed by extruding the circle across cortical depth. We also calculated total length and density at different cortical depth. Specifically, we calculated total length and density with 10 μm step from 0 – 600 μm to generate the depth profile. We also calculated mean density at different depth range to compare the layer specificity (superficial layer: 0 – 150 μm; middle layer: 150 - 350 μm; deep layer: 350 - 600 μm).

### Statistics

The imaging preprocessing, nwb packaging, bouton clustering, receptive field analysis, grating response analysis, functional type classification, and axon morphology analysis were performed by a custom-written python package “NeuroAnalysisTools” (https://github.com/zhuangjun1981/NeuroAnalysisTools).

To control the variability across imaging planes (different axon/bouton density, vasculature pattern, expression level), most statistics were extracted from each plane and separated for different functional types if necessary. For statistics that only one number can be drawn from each imaging plane (e.g. bouton count), they were presented as mean ± standard deviation. For statistics that a population distribution can be drawn from each imaging plane (gDSI, gOSI, peak SF/TF, RF strength, bouton per cluster, max bouton distance, axon coverage area), mean for each plane was calculated first and mean ± s.e.m. was reported across imaging planes. The comparisons between different functional types within imaging plane were performed by Wilcoxon rank-sum test, and comparisons between imaging planes were performed by Mann-Whitney U test, if not otherwise stated. Nonetheless, we have verified our results of these two tests by paired t-test and independent t-test respectively, and they all agreed with the non-parametric tests (not shown).

## Acknowledgments

Research reported in this publication was supported by the National Institute of Neurological Disorders and Stroke (Award Number R01NS104949) and the National Institute of Mental Health (Award Number R01MH117820) of the National Institutes of Health. The content is solely the responsibility of the authors and does not necessarily represent the official views of the National Institutes of Health. We thank Dr. Saskia de Vries, Dr. Douglas Storace, Dr. Pooja Balaram, and Dr. Daniel Millman for their comments and suggestions. We thank Dr. Roger Tsien for providing tdTomato construct. We thank many staff members of the Allen Institute, especially the In Vivo Sciences team for surgeries and the Manufacturing and Processing Engineering team for hardware support. We wish to thank the Allen Institute for Brain Science founder, Paul G. Allen, for his vision, encouragement, and support.

---

**Figure S1.**
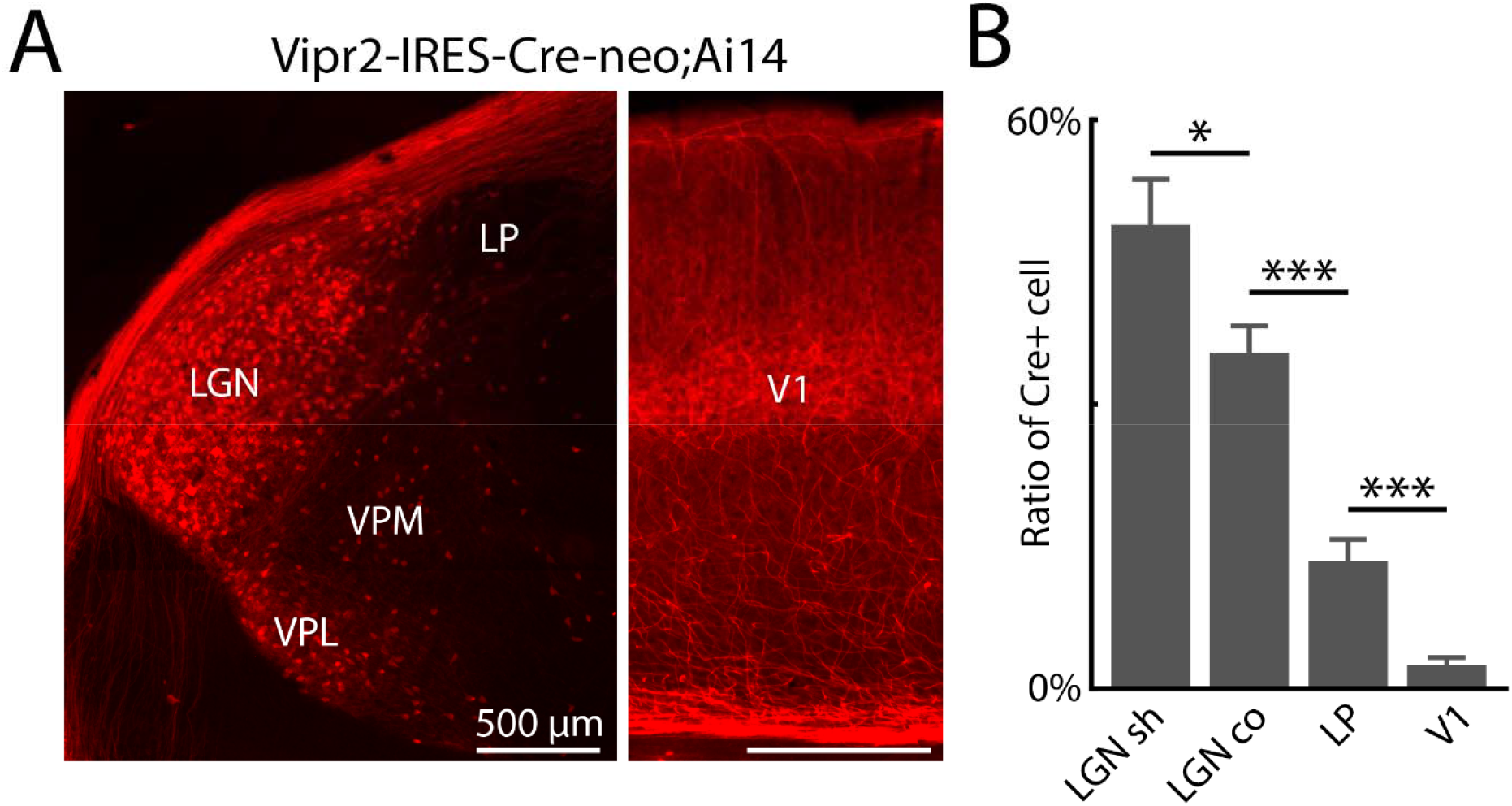
Cre expression characterization of Vipr2-IRES-Cre-neo mouse line. **(A)** Coronal sections with tdTomato positive cells in thalamus (left) and axons in V1 (right) from a Vipr2-IRES-Cre-neo;Ai14 mouse. **(B)** Ratio of Cre+ cells to Nissl labeled cells in visual thalamus and V1. Data from Allen Brain Atlas Transgenic Characterization dataset (experiments: <u>576523754</u>, <u>576524006</u>). 10 fields of view were manually selected for each brain region. For each field of view, Cre+ cells and Nissl+ cells were manually counted and the ratio between the two are calculated. dLGN sh: dLGN shell. dLGN co: dLGN core. LP: lateral posterior nucleus of thalamus. V1: primary visual cortex. Bar graph: mean ± sem. Independent t-test: dLGN sh vs. dLGN co: t=2.3, p=0.034; dLGN co vs. LP, t=5.8, p=1.8×10^−5^; LP vs. V1, t=4.4, p=0.00037

**Figure S2.**
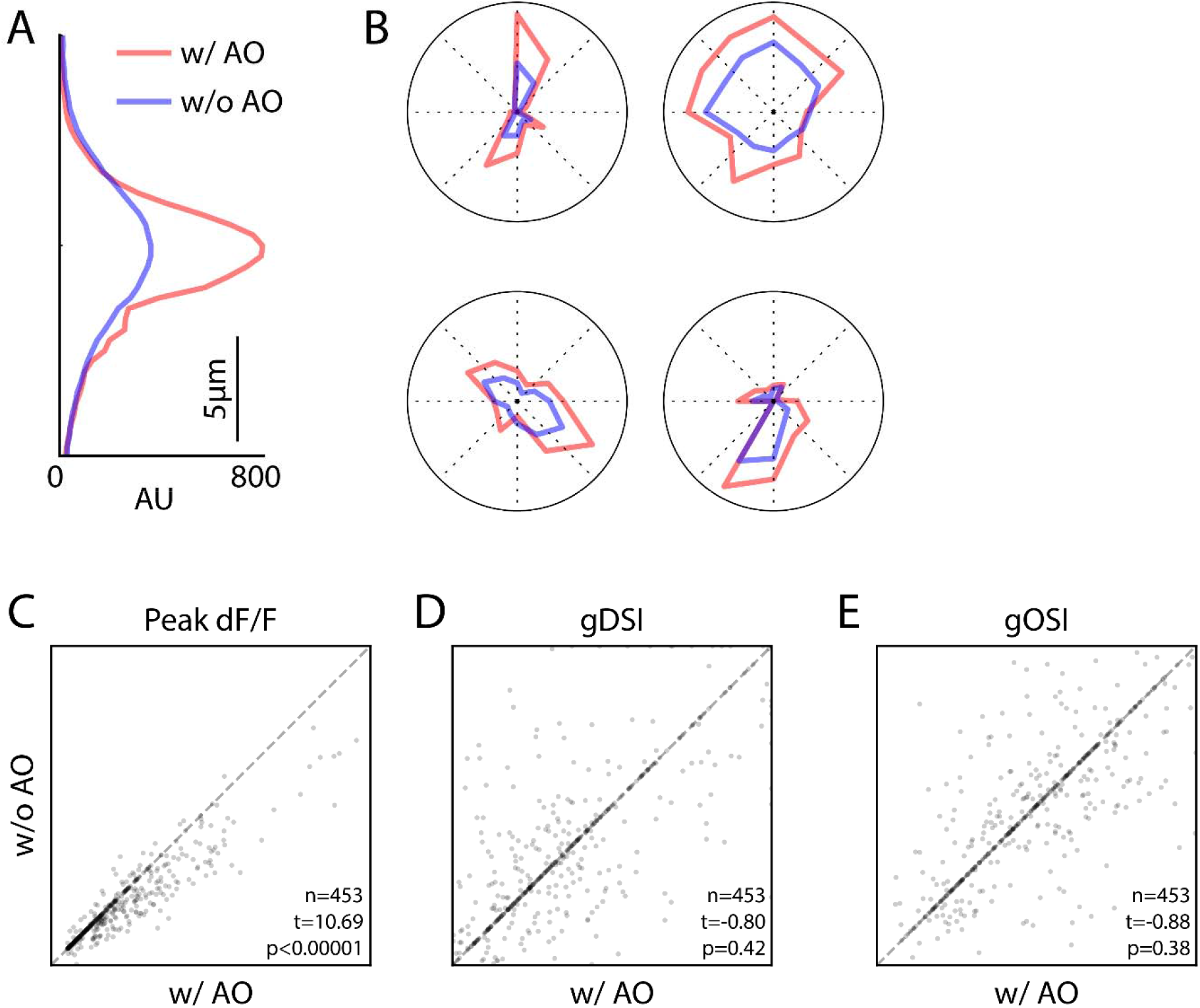
Comparisons of response amplitude, direction selectivity and orientation selectivity between conditions with and without adaptive optics (AO). **(A)** Point spread function with and without AO by imaging 2 μm beads deposited between a coverglass and the brain surface in an awake mouse. **(B)** Example orientation/direction tuning curves from 4 example boutons. **(C - E)** Comparisons of peak dF/F, gDSI, and gOSI, respectively, between with AO and without AO conditions. Two imaging planes in two mice (300 μm and 350 μm below pia). In AO conditions, laser beam wave front was corrected separately for each mouse. Significance test: paired t-test. Statistics are shown as insets.

**Figure S3.**
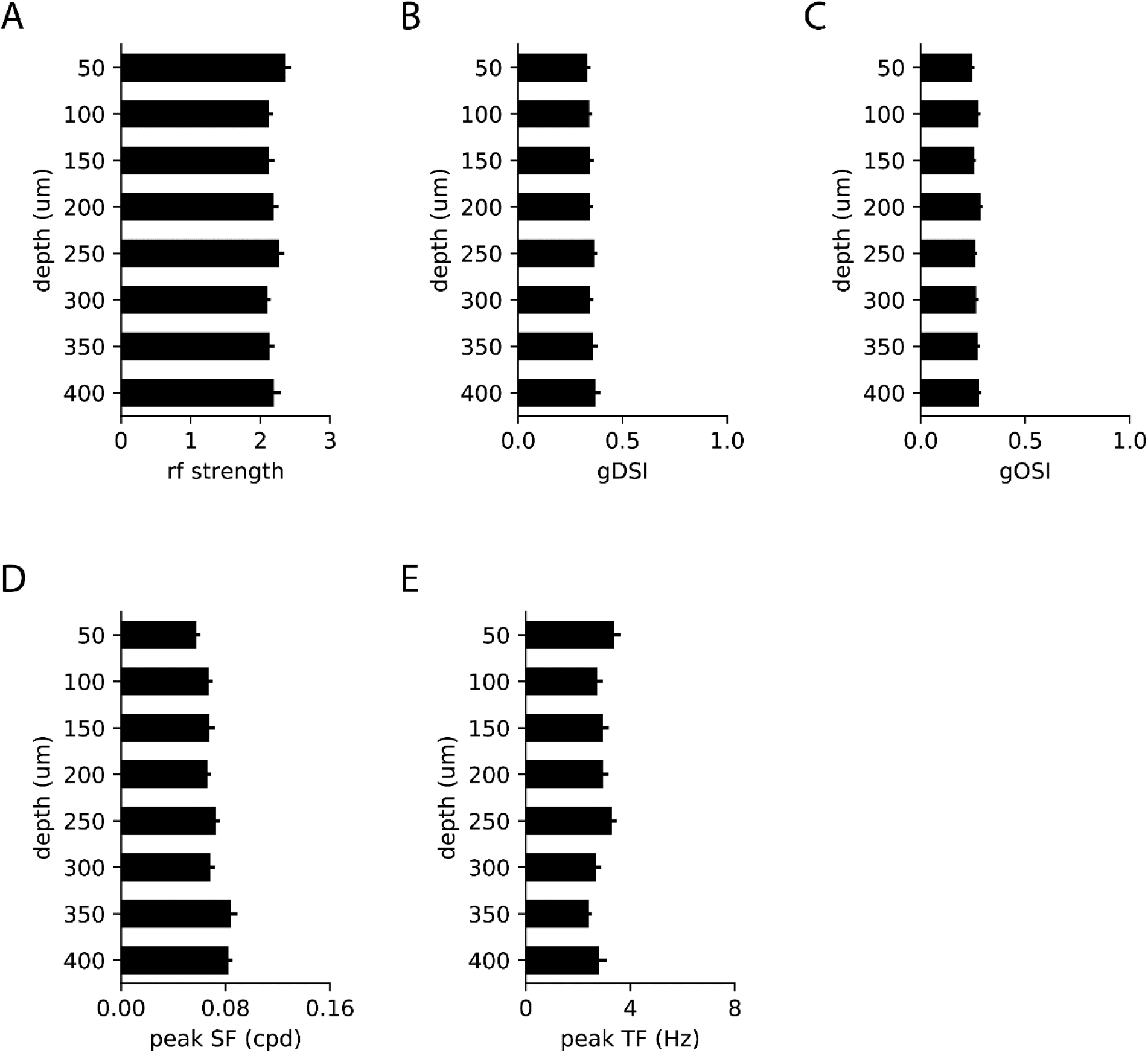
Metrics across depth. **(A - E)** different response properties across cortical depth. For each metric, the mean of each imaging plane was calculated first and the mean and s.e.m. across depths were plotted here. For gDSI, gOSI, peak SF, and peak TF, only boutons with significant responses to drifting gratings (Methods) were included. One-way ANOVA, RF strength: F=1.47, p=0.19; gDSI: F=0.42, p=0.89; gOSI: F=1.76; p=0.10; peak SF: F=3.79, p=0.001; peak TF: F=1.70, p=0.12.

**Figure S4.**
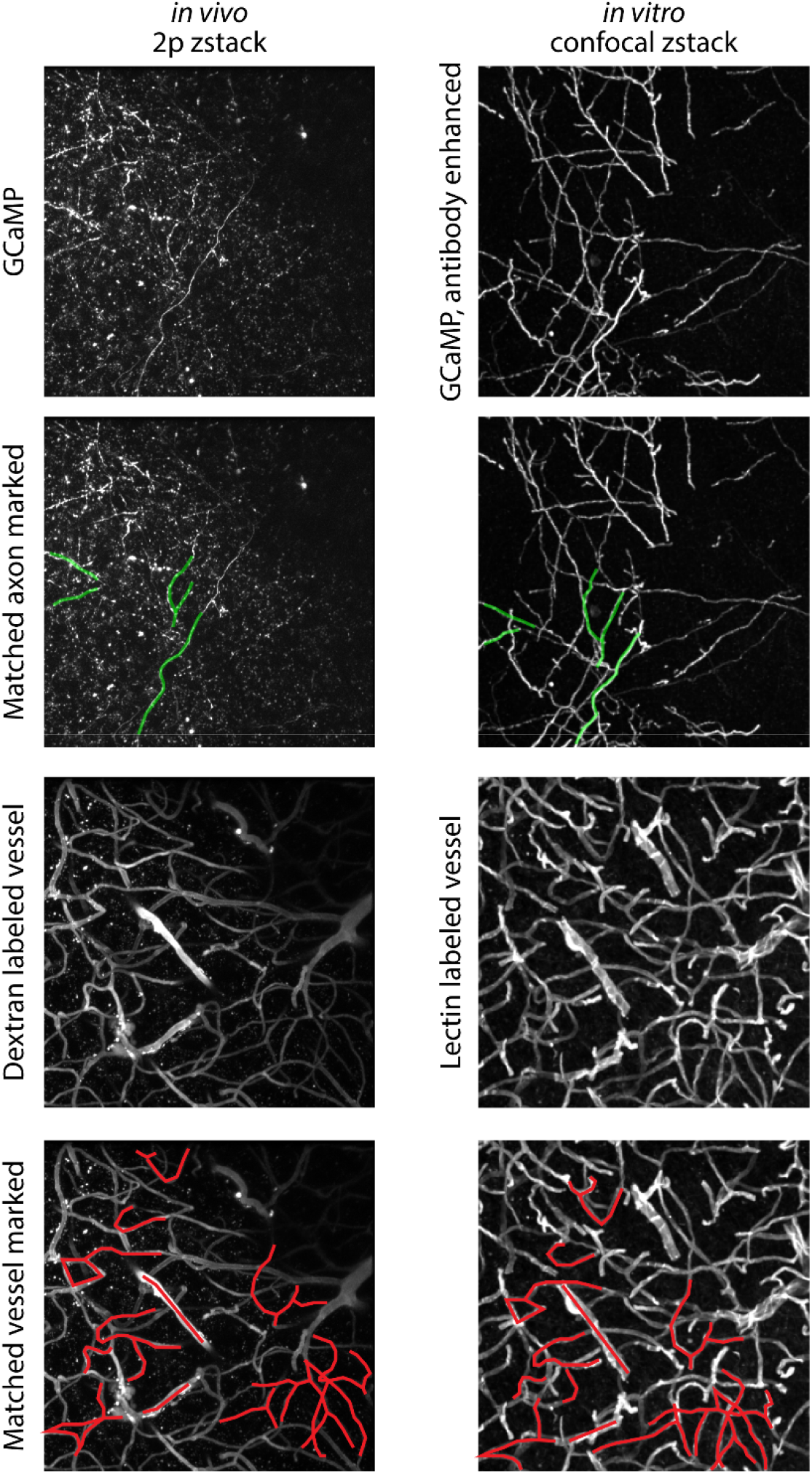
Coregistration between *in vivo* two-photon volumes and *in vitro* confocal volumes. For *in vivo* two-photon imaging (left column), the dLGN axons were sparsely labeled with GCaMP by AAV viral labeling and blood vessels were labeled by fluorescence dye (dextran-Texas Red). For *in vitro* confocal imaging (right column), the imaged tissue volume was collected and cleared, the GCaMP signal was enhanced by antibody labeling, and blood vessels were labeled by fluorescence dye (lectin-DyLight). Blood vessels labeled in both imaging modalities were used as fiducial structure for locating the same region in both two-photon and confocal images (bottom row, red lines), and the same axons imaged *in vivo* were identified in the confocal volumes (second row, green lines).

## Notes

### Competing Interest Statement

The authors have declared no competing interest.

## References

Anderson JC, da Costa NM, Martin KA (2009) The W cell pathway to cat primary visual cortex. J Comp Neurol 516:20–35.

Antonini A, Fagiolini M, Stryker MP (1999) Anatomical correlates of functional plasticity in mouse visual cortex. J Neurosci 19:4388–4406.

Bereshpolova Y, Stoelzel CR, Su C, Alonso JM, Swadlow HA (2019) Activation of a Visual Cortical Column by a Directionally Selective Thalamocortical Neuron. Cell Rep 27:3733–3740 e3733.

Cruz-Martin A, El-Danaf RN, Osakada F, Sriram B, Dhande OS, Nguyen PL, Callaway EM, Ghosh A, Huberman AD (2014) A dedicated circuit links direction-selective retinal ganglion cells to the primary visual cortex. Nature 507:358–361.

Durand S, Iyer R, Mizuseki K, de Vries S, Mihalas S, Reid RC (2016) A Comparison of Visual Response Properties in the Lateral Geniculate Nucleus and Primary Visual Cortex of Awake and Anesthetized Mice. J Neurosci 36:12144–12156.

Denman, D.J., and Contreras, D. (2016). On Parallel Streams through the Mouse Dorsal Lateral Geniculate Nucleus. Front. Neural Circuits 10, 20.

Grubb MS, Thompson ID (2003) Quantitative characterization of visual response properties in the mouse dorsal lateral geniculate nucleus. J Neurophysiol 90:3594–3607.

Hei X, Stoelzel CR, Zhuang J, Bereshpolova Y, Huff JM, Alonso JM, Swadlow HA (2014) Directional selective neurons in the awake dLGN: response properties and modulation by brain state. J Neurophysiol 112:362–373.

Kondo S, Ohki K (2016) Laminar differences in the orientation selectivity of geniculate afferents in mouse primary visual cortex. Nat Neurosci 19:316–319.

Liang L, Fratzl A, Goldey G, Ramesh RN, Sugden AU, Morgan JL, Chen C, Andermann ML (2018) A Fine-Scale Functional Logic to Convergence from Retina to Thalamus. Cell 173:1343–1355.

Lien, A.D., and Scanziani, M. (2013). Tuned thalamic excitation is amplified by visual cortical circuits. Nat Neurosci 16, 1315–1323.

Marshel JH, Kaye AP, Nauhaus I, Callaway EM (2012) Anterior-posterior direction opponency in the superficial mouse lateral geniculate nucleus. Neuron 76:713–720.

Peng H, Xie P, Liu L, et al. (2020) Brain-wide single neuron reconstruction reveals morphological diversity in molecularly defined striatal, thalamic, cortical and claustral neuron types. bioRxiv 675280.

Piscopo DM, El-Danaf RN, Huberman AD, Niell CM (2013) Diverse visual features encoded in mouse lateral geniculate nucleus. J Neurosci 33:4642–4656.

Román Rosón, M., Bauer, Y., Kotkat, A.H., Berens, P., Euler, T., and Busse, L. (2019). Mouse dLGN Receives Functional Input from a Diverse Population of Retinal Ganglion Cells with Limited Convergence. Neuron 102, 462–476.e8.

Seabrook TA, Burbridge TJ, Crair MC, Huberman AD (2017) Architecture, Function, and Assembly of the Mouse Visual System. Annu Rev Neurosci 40:499–538.

Sun W, Tan Z, Mensh BD, Ji N (2016) Thalamus provides layer 4 of primary visual cortex with orientation- and direction-tuned inputs. Nat Neurosci 19:308–315.

Susaki, E.A., Tainaka, K., Perrin, D., Yukinaga, H., Kuno, A., and Ueda, H.R. (2015). Advanced CUBIC protocols for whole-brain and whole-body clearing and imaging. Nat. Protoc. 10, 1709–1727.

Sur M, Sherman SM (1982) Linear and nonlinear W-cells in C-laminae of the cat’s lateral geniculate nucleus. J Neurophysiol 47:869–884.

Suresh, V., Ciftcioglu, U.M., Wang, X., Lala, B.M., Ding, K.R., Smith, W.A., Sommer, F.T., and Hirsch, J.A. (2016). Synaptic Contributions to Receptive Field Structure and Response Properties in the Rodent Lateral Geniculate Nucleus of the Thalamus. J Neurosci 36, 10949–10963.

Zhao X, Chen H, Liu X, Cang J (2013) Orientation-selective responses in the mouse lateral geniculate nucleus. J Neurosci 33:12751–12763.

## Additional references for Methods

Bria, A., and Iannello, G. (2012). TeraStitcher - A tool for fast automatic 3D-stitching of teravoxel-sized microscopy images. BMC Bioinformatics 13.

de Vries, S.E.J., Lecoq, J.A., Buice, M.A. et al. (2020) A large-scale standardized physiological survey reveals functional organization of the mouse visual cortex. Nat Neurosci 23:138–151

Dipoppa M, Ranson A, Krumin M, Pachitariu M, Carandini M, Harris KD (2018) Vision and Locomotion Shape the Interactions between Neuron Types in Mouse Visual Cortex. Neuron 98:602–615 e608.

Economo, M.N., Clack, N.G., Lavis, L.D., Gerfen, C.R., Svoboda, K., Myers, E.W., and Chandrashekar, J. (2016). A platform for brain-wide imaging and reconstruction of individual neurons. Elife 5, e10566.

Giovannucci A, Friedrich J, Gunn P, Kalfon J, Brown BL, Koay SA, Taxidis J, Najafi F, Gauthier JL, Zhou P, Khakh BS, Tank DW, Chklovskii DB, Pnevmatikakis EA (2019) CaImAn an open source tool for scalable calcium imaging data analysis. Elife 8:e38173.

Goldey GJ, Roumis DK, Glickfeld LL, Kerlin AM, Reid RC, Bonin V, Schafer DP, Andermann ML (2014) Removable cranial windows for long-term imaging in awake mice. Nat Protoc 9:2515–2538.

Juavinett AL, Nauhaus I, Garrett ME, Zhuang J, Callaway EM (2017) Automated identification of mouse visual areas with intrinsic signal imaging. Nat Protoc 12:32–43.

Liu, R., Ball, N., Brockill, J., Kuan, L., Millman, D., White, C., Leon, A., Williams, D., Nishiwaki, S., de Vries, S., et al. (2019). Aberration-free multi-plane imaging of neural activity from the mammalian brain using a fast-switching liquid crystal spatial light modulator. Biomed. Opt. Express 10, 5059.

Madisen, L., Zwingman, T.A., Sunkin, S.M., Oh, S.W., Zariwala, H.A., Gu, H., Ng, L.L., Palmiter, R.D., Hawrylycz, M.J., Jones, A.R., et al. (2010). A robust and high-throughput Cre reporting and characterization system for the whole mouse brain. Nat Neurosci 13, 133–140.

Mukamel EA, Nimmerjahn A, Schnitzer MJ (2009) Automated analysis of cellular signals from large-scale calcium imaging data. Neuron 63:747–760.

Peirce J, Gray JR, Simpson S, MacAskill M, Hochenberger R, Sogo H, Kastman E, Lindelov JK (2019) PsychoPy2: Experiments in behavior made easy. Behav Res Methods 51:195–203.

Petreanu, L., Gutnisky, D.A., Huber, D., Xu, N.L., O’Connor, D.H., Tian, L., Looger, L., and Svoboda, K. (2012). Activity in motor-sensory projections reveals distributed coding in somatosensation. Nature 489, 299–303.

Pnevmatikakis EA, Soudry D, Gao Y, Machado TA, Merel J, Pfau D, Reardon T, Mu Y, Lacefield C, Yang W, Ahrens M, Bruno R, Jessell TM, Peterka DS, Yuste R, Paninski L (2016) Simultaneous Denoising, Deconvolution, and Demixing of Calcium Imaging Data. Neuron 89:285–299.

Pologruto TA, Sabatini BL, Svoboda K (2003) ScanImage: flexible software for operating laser scanning microscopes. Biomed Eng Online 2:13.

Zhuang J, Stoelzel CR, Bereshpolova Y, Huff JM, Hei X, Alonso JM, Swadlow HA (2013) Layer 4 in primary visual cortex of the awake rabbit: contrasting properties of simple cells and putative feedforward inhibitory intercs. J Neurosci 33:11372–11389.

Zhuang J, Ng L, Williams D, Valley M, Li Y, Garrett M, Waters J (2017) An extended retinotopic map of mouse cortex. Elife 6:e18372.

